# A Centimeter-Scale, Peristaltic Human Intestinal Organoid with Integrated Neuro-Immune-Vascular Systems Recapitulates Enteritis and Orthotopic Colorectal Cancer

**DOI:** 10.64898/2026.08.18.745659

**Authors:** Zhen Qi, Min Shen, Kang Wang, Xiaowen Li, Mingqian Huang, Yandan Liu, Yue yu, Zongjun Liu

## Abstract

Human pluripotent stem cell-derived intestinal organoids hold great promise for disease modeling, drug screening, and regenerative medicine. However, conventional intestinal organoids are predominantly epithelial, small in scale, and lack the multicellular complexity required to recapitulate the pathophysiology of intestinal disorders such as inflammatory bowel disease (IBD) and colorectal cancer (CRC). Here, we report the development of Centimeter-Scale, purely 3D self-organized human intestinal organoids (IOs) from induced pluripotent stem cells (iPSCs) that encompass multiple tissue lineages, including epithelium, mesenchyme, smooth muscle, neurons, immune cells, and vasculature. These organoids achieve functional maturation by day 100+, exhibiting rhythmic peristaltic-like contractions, and by day 147 they display histological structures including lumens, crypt-like architecture, goblet cells, and smooth muscle. Importantly, for the first time, the neuroOmuscle lineages arise spontaneously and autonomously in a purely 3D culture system, without any external stimulation (e.g., electrical, chemical, or mechanical), and mature to form functional neuromuscular junctions, driving macroscopically visible peristalticOlike contractions that mimic intestinal motility entirely through in vitro culture, without any xenotransplantation. Single-cell RNA sequencing at day 115 identified 12 cell subtypes across four major lineages, recapitulating the cellular diversity of the developing human intestine. Using this platform, we established an LPS/IFN-γ-induced IBD model that recapitulated key pathological features, including epithelial disruption, immune cell infiltration, and IL-6 elevation. Transcriptomic analysis confirmed activation of the NF-κB and JAK2-STAT3 pathways, multi-modal cell death, and immune recruitment machinery — all consistent with clinical IBD pathology. Furthermore, we developed intestinal cancer models at 7 and 21 days showing abnormal hyperplasia, and a probiotic co-culture system demonstrating anti-inflammatory efficacy. Together, these results establish Centimeter-Scale intestinal organoids as a physiologically relevant, multicellular platform for modeling intestinal diseases and evaluating therapeutic interventions.

## Introduction

The human intestine is a complex organ composed of multiple interacting tissue compartments, including the epithelial lining, lamina propria mesenchyme, smooth muscle layers, enteric nervous system (ENS), and mucosal immune cells. The coordinated function of these compartments is essential for nutrient absorption, barrier defense, immune homeostasis, and motility. Disruptions to this delicate balance underlie prevalent intestinal disorders, including inflammatory bowel disease (IBD) — encompassing ulcerative colitis (UC) and Crohn’s disease (CD) — and colorectal cancer (CRC). Together, these disorders affect tens of millions of individuals worldwide and represent a major unmet medical need (Ng et al., 2018; Sung et al., 2021).

Intestinal organoids have emerged as powerful in vitro models that bridge the gap between simplified cell line cultures and animal models. Pioneered by Clevers and colleagues using Lgr5+ intestinal stem cells (Sato et al., 2009), organoid technology has since been extended to human pluripotent stem cells (hPSCs), enabling the derivation of intestinal tissues from iPSCs through directed differentiation protocols that mimic embryonic development (Spence et al., 2011; Watson et al., 2014). Despite significant advances, most current intestinal organoid models remain limited in several critical aspects. First, conventional organoids are typically small (<500 μm) and cultured in Matrigel domes, restricting their utility for modeling spatial tissue architecture. Second, they are predominantly epithelial, lacking the mesenchymal, immune, neuronal, and vascular compartments essential for recapitulating the multi-cellular pathophysiology of IBD and CRC. Third, functional maturation — particularly the emergence of motility — has been difficult to achieve consistently.

To address these limitations, we developed a purely 3D self-organizing differentiation protocol that generates centimeter-scale human intestinal organoids from iPSCs. Unlike air-liquid interface (ALI) cultures or co-culture approaches that require exogenous stromal components, our method relies entirely on the intrinsic self-organization capacity of iPSC-derived progenitors, yielding organoids that spontaneously develop multiple tissue lineages. The entire differentiation proceeds in a purely 3D format that is straightforward to perform with high success rates, and it is completely free of Matrigel, making the system directly compatible with bioreactor scale-up for high-throughput production under defined, stable, and fully controllable culture conditions. These organoids achieve advanced functional maturation, as evidenced by rhythmic peristaltic-like contractions at day 100+, and histological complexity by day 147, including lumens, crypt-like structures, goblet cells, and smooth muscle. Notably, the neuro-muscle lineages arise autonomously in vitro, mature to form functional neuromuscular junctions, and drive macroscopically visible peristaltic-like contractions without any transplantation into animals; the resulting organoids represent the largest reported size for an intestinal organoid and remain robust and motile for over 120 days in culture.

In this study, we systematically characterized these Centimeter-Scale intestinal organoids using histology, immunofluorescence, and single-cell RNA sequencing. We then applied this platform to model three clinically relevant scenarios: (1) IBD, using LPS and IFN-γ stimulation to recapitulate inflammatory epithelial damage; (2) CRC, using a chemical induction approach at 7 and 21 days; and (3) probiotic-host interactions, using a fluorescence-labeled probiotic injection system. Our results demonstrate that centimeter-scale, multicellular intestinal organoids provide a physiologically relevant platform for intestinal disease modeling, offering advantages over conventional epithelial-only organoids for studying multi-tissue pathogenesis and evaluating therapeutic candidates.

## Results

### Generation and Functional Maturation of Centimeter-Scale Intestinal Organoids

We developed a directed differentiation protocol to generate Centimeter-Scale intestinal organoids from human iPSCs (Figure 1). The differentiation timeline spans from iPSC maintenance (day −3) through definitive endoderm induction and intestinal specification (day 0–9), followed by long-term maintenance in a proprietary intestinal organoid maintenance medium (MIDIO-001). By day 9, three-dimensional organoid structures became visible, which continued to expand in scale over the following weeks. By day 30, organoids reached sufficient size for drug screening and disease model construction. Notably, by day 100+, organoids exhibited rhythmic peristaltic-like contractions — a hallmark of advanced functional maturation that recapitulates the motility of the native intestine (Figure 1). These contractions, observed as spontaneous, periodic waves of muscular compression propagating along the organoid body, indicate the establishment of functional smooth muscle-neuron units within the organoids. Importantly, these contractions were visible to the naked eye and arose entirely in vitro — without any transplantation into mice — demonstrating autonomous generation of the neuro-muscle lineage, its maturation in vitro, and the formation of functional neuromuscular connections within the organoids.

**Figure 1.**
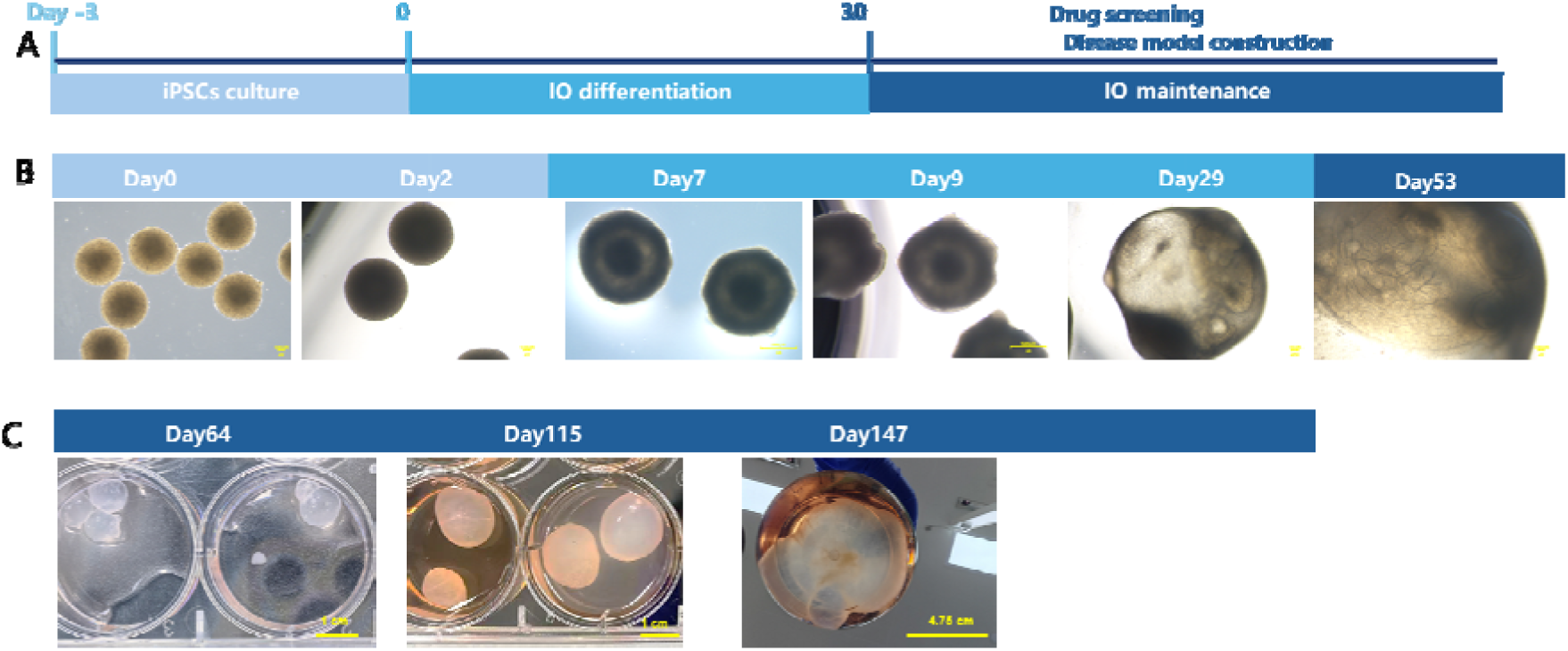
Generation and functional maturation of Centimeter-Scale intestinal organoids. (A) Schematic overview of the intestinal organoid differentiation protocol, showing key stages from iPSC culture (day −3) through differentiation (day 0–9), maintenance (day 9–53), and functional maturation (day 100+). (B) Brightfield images of intestinal organoids at days 0, 2, 7, 9, 29 and 53 showing progressive growth and morphological complexity. Scale bars: 500 μm. (C) Representative images showing intestinal organoids enlarging from Day 64 to 147. Initially round, they develop buds and lumens by Day 100, becoming centimeter-scale with complex morphology and thickened walls at Day 115–147. Scale bars: 1 cm.

The organoids continued to grow and mature in long-term culture, reaching centimeter-scale dimensions. At these dimensions, they represent, to our knowledge, the largest reported size for an intestinal organoid. The sustained growth and functional maturation were maintained using the proprietary maintenance medium (MIDIO-001), which supports the long-term viability and multilineage preservation of the organoids. Strikingly, the organoids remained viable and peristaltic for over 120 days of continuous culture, underscoring the exceptional robustness and long-term stability of the platform.

### Intestinal Organoids Recapitulate the Architecture of the Human Intestine

To assess the histological complexity of the organoids, we performed H&E staining on day 147 intestinal organoids. The analysis revealed multiple tissue structures characteristic of the human intestinal wall (Figure 2). Organoids displayed well-defined lumens lined by epithelial cells, intestinal crypt-like invaginations extending into the surrounding mesenchyme, mature goblet cells containing mucin-filled vacuoles, and organized smooth muscle-like cell layers. These findings demonstrate that the 3D self-organization process yields tissue architecture that closely resembles the native intestinal organization, rather than the disorganized cell aggregates typical of conventional organoid cultures.

**Figure 2.**
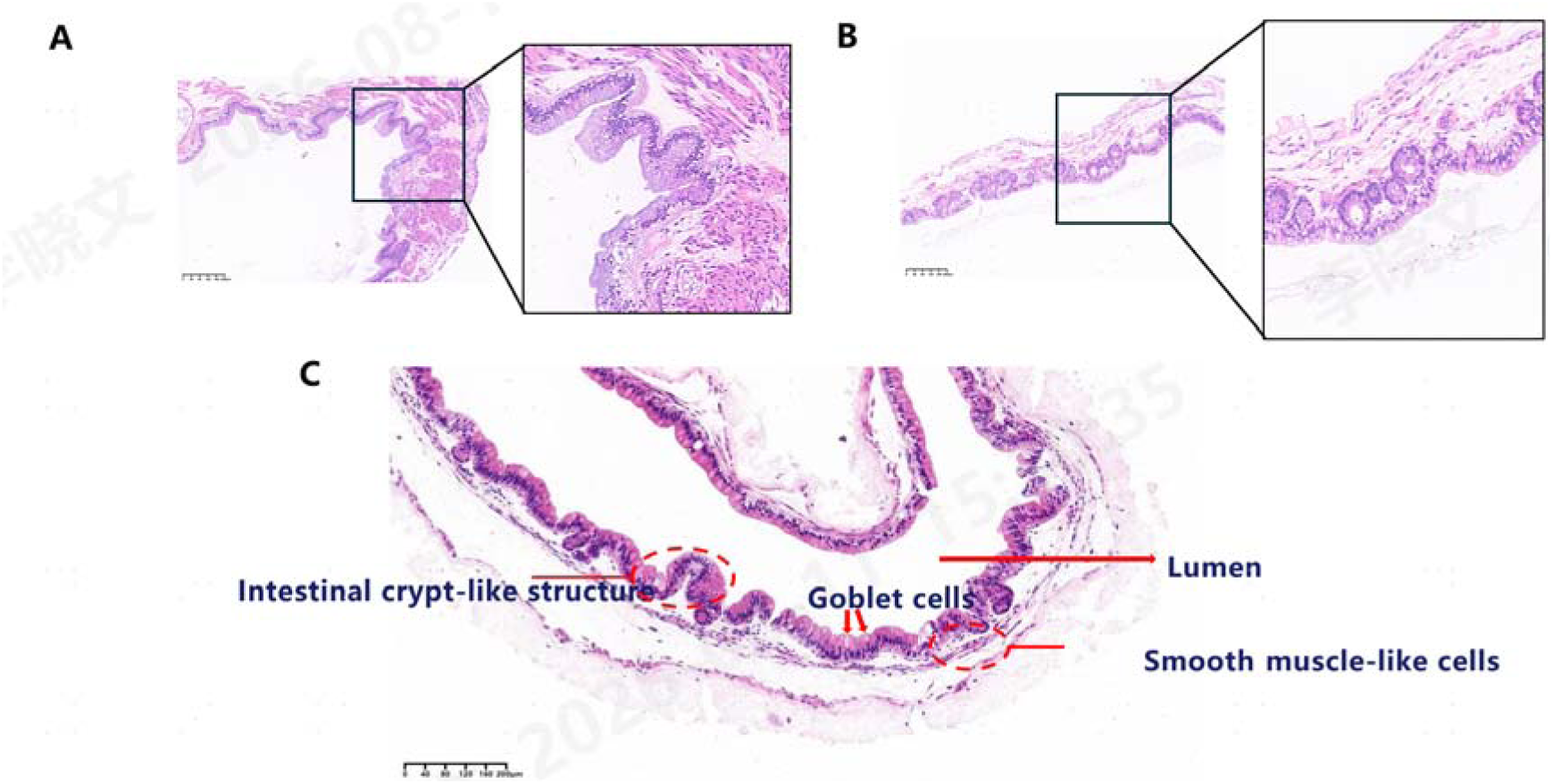
Intestinal organoids recapitulate the architecture of the human intestine. (A) H&E staining of a day 64 intestinal organoid showing overall tissue organization. Scale bar: 200 μm. (C) H&E staining of a day 147 intestinal organoid showing lumen lined by epithelial cells, intestinal crypt-like invaginations, goblet cells with mucin vacuoles, and (E) smooth muscle-like cell layers. Scale bars: 50 μm.

### Multilineage Cell Type Diversity Revealed by Immunofluorescence

To further characterize the cellular composition of the organoids, we performed multiplex immunofluorescence staining on day 64 intestinal organoids (Figure 3). Staining for EPCAM confirmed the presence of intestinal epithelial cells, which formed continuous layers lining the organoid surfaces and lumens. We identified CD68+ immune cells, including macrophage-like cells, within the organoid stroma, demonstrating the presence of endogenous immune populations without exogenous addition. CHGA+ neuroendocrine cells were interspersed within the epithelial layer, indicating the maturation of enteroendocrine lineages. α-SMA+ We found smooth muscle cells organized in bands surrounding the epithelial structures, consistent with the formation of muscular layers. TUJ1+ neurons appeared in proximity to the smooth muscle, suggesting the development of enteric nervous system (ENS) components. Furthermore, whole-mount immunofluorescence staining of day 100+ organoids revealed CD31+ endothelial cells forming vascular-like networks alongside EPCAM+ epithelial structures, indicating the emergence of rudimentary vasculature within the organoids (Figure 3).

**Figure 3.**
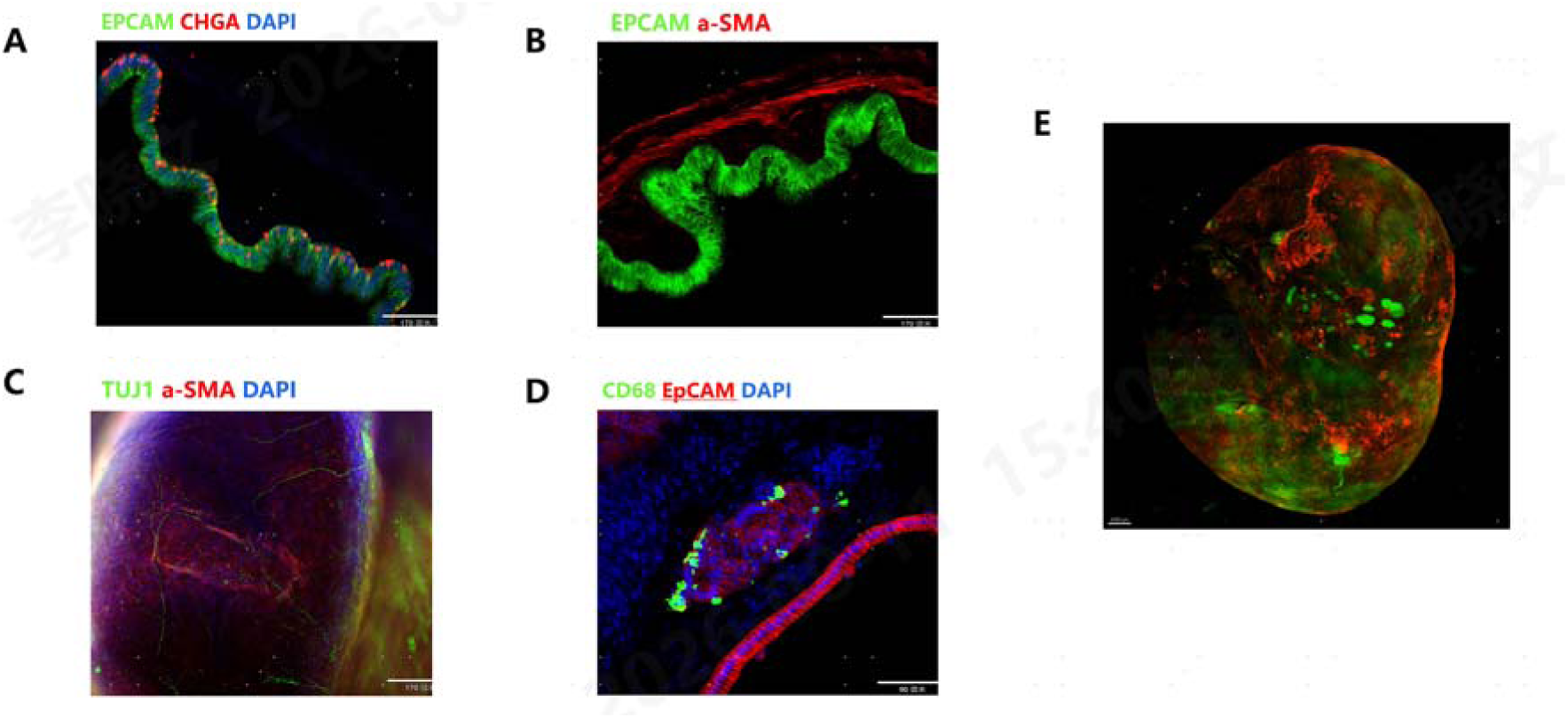
Multilineage cell type diversity in intestinal organoids. (A–D) Immunofluorescence staining of day 64 intestinal organoid sections: (A) EPCAM and CHGA (neuroendocrine cells). Scale bars: 170 μm. (B) EPCAM and α-SMA (smooth muscle cells). Scale bars: 170 μm. (C) TUJ1 (neurons) and α-SMA. Scale bars: 170 μm. (D) EPCAM (epithelial cells) and CD68 (immune cells). Scale bars: 90 μm. Nuclei counterstained with DAPI. Scale bars: 90 μm. (E) Whole-mount immunofluorescence of day 100+ organoid showing CD31+ vascular structures (green) and EPCAM+ epithelium (red). Scale bar: 100 μm.

### Single-Cell Transcriptomic Profiling Reveals Cellular Heterogeneity

To comprehensively map the cellular diversity of the organoids, we performed single-cell RNA sequencing (scRNA-seq) on day 115 intestinal organoids. After quality filtering, we retained 7,960 cells for analysis. Unsupervised clustering and cell type annotation based on canonical marker genes identified 12 distinct cell subtypes spanning four major lineages (Figure 4, Table 1):

**Figure 4.**
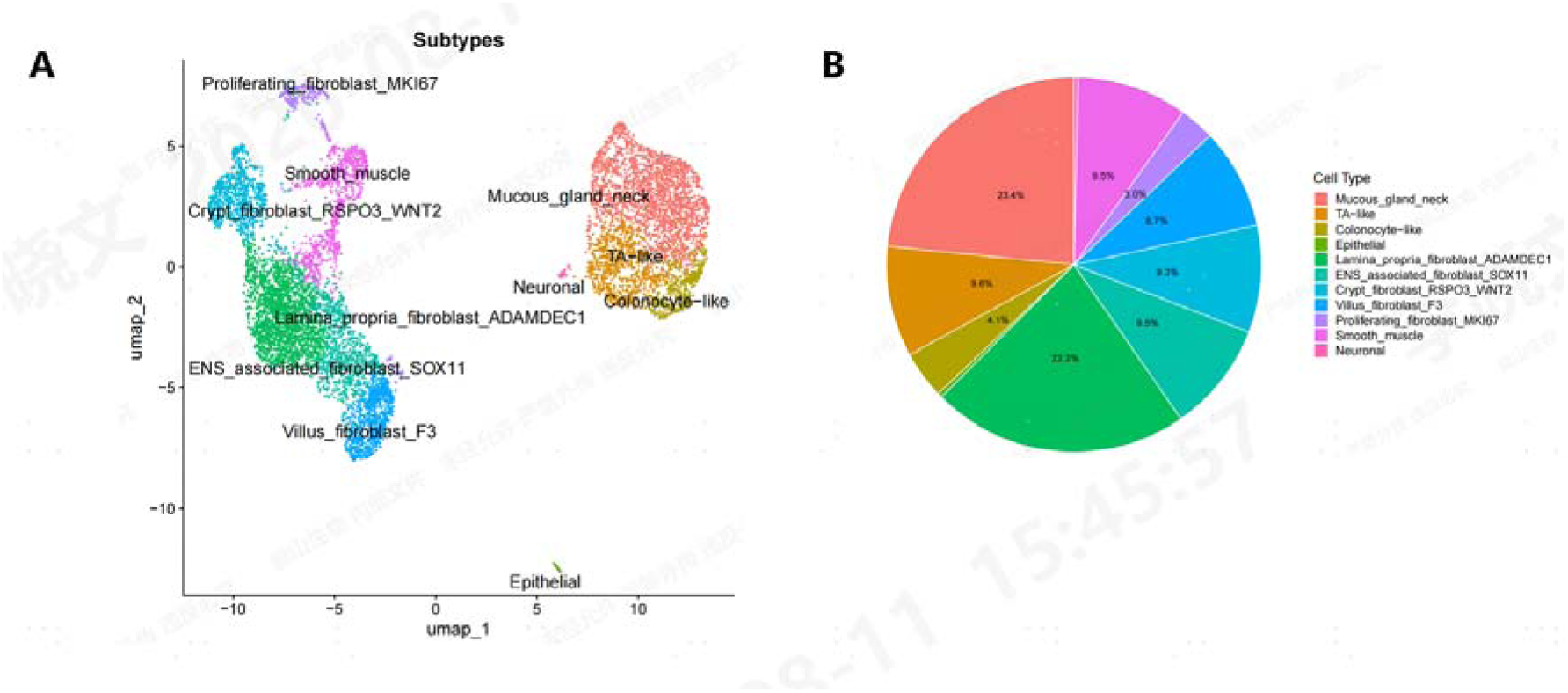
Single-cell transcriptomic profiling of day 115 intestinal organoids. (A) UMAP visualization of 7,960 cells, colored by annotated cell subtype (12 subtypes across four lineages: epithelial, mesenchymal, smooth muscle, and neuronal). (B) Pie chart showing the proportion of each cell subtype. Epi: epithelial; Mes: mesenchymal; SM: smooth muscle; Neu: neuronal.

**Table 1.**
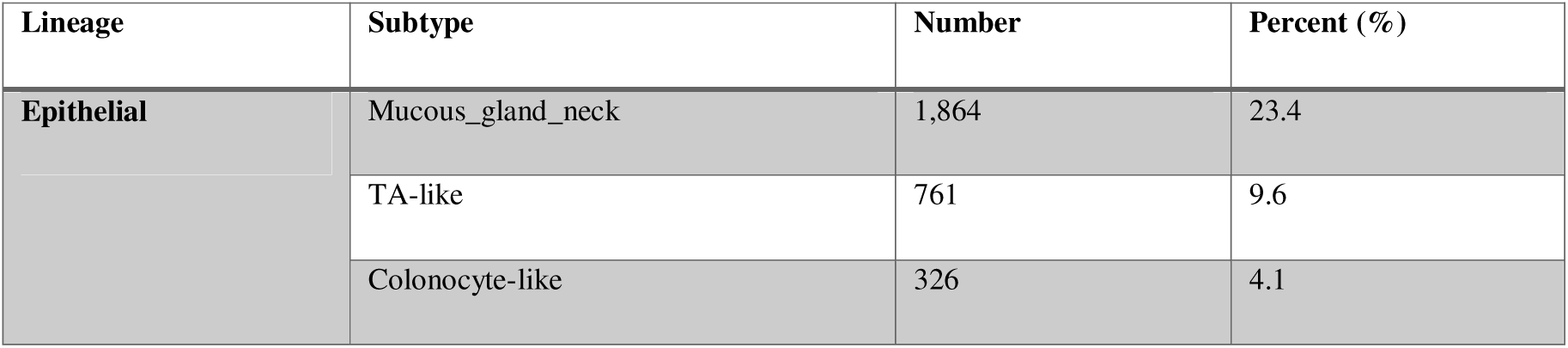
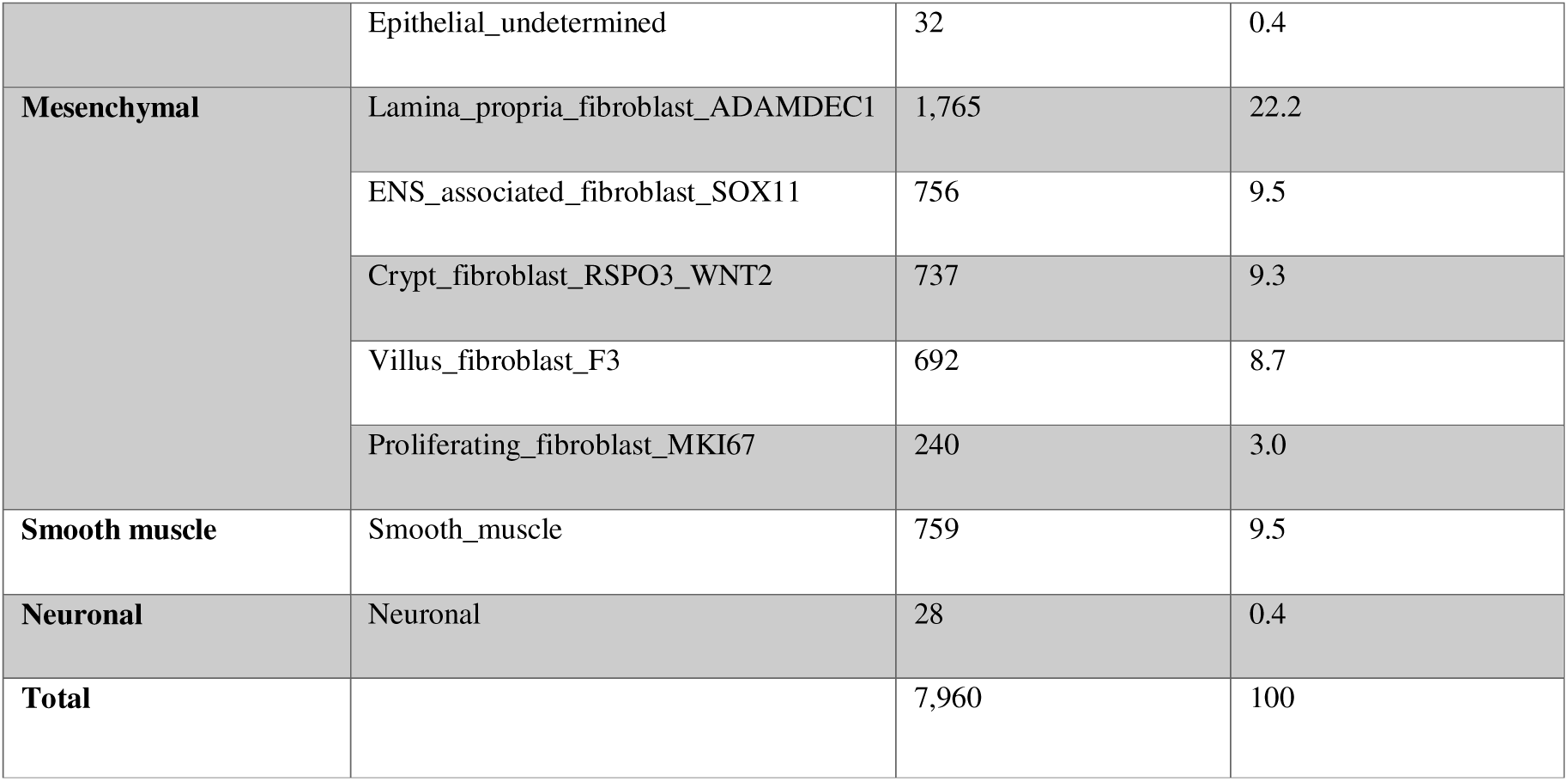
Single-cell RNA-seq cell subtype composition of day 115 intestinal organoids.

Epithelial lineage (37.5%): The largest epithelial population was mucous/gland neck cells (23.4%), expressing mucin genes (MUC2, TFF3), followed by transit-amplifying (TA)-like cells (9.6%) expressing proliferation markers (MKI67, PCNA), colonocyte-like cells (4.1%) expressing absorptive markers, and a small population of undetermined epithelial cells (0.4%).

Mesenchymal lineage (52.7%): The most abundant mesenchymal subtype was lamina propria fibroblasts expressing ADAMDEC1 (22.2%), followed by ENS-associated fibroblasts expressing SOX11 (9.5%), crypt fibroblasts expressing RSPO3 and WNT2 (9.3%), villus fibroblasts expressing F3 (8.7%), and proliferating fibroblasts expressing MKI67 (3.0%). The presence of multiple fibroblast subtypes with distinct signaling roles (e.g., WNT2/RSPO3 from crypt fibroblasts supporting epithelial stemness) indicates a well-organized mesenchymal niche.

Smooth muscle lineage (9.5%): A distinct cluster of smooth muscle cells expressing ACTA2, MYH11, and DESMIN was identified, consistent with the α-SMA+ cells observed by immunofluorescence.

Neuronal lineage (0.4%): A small but detectable population of neuronal cells expressing neuronal markers was identified, confirming the presence of ENS-derived cells within the organoids. These results demonstrate that the Centimeter-Scale intestinal organoids encompass the major cell types of the intestinal wall, recapitulating the development, architecture, and function of the fetal intestine in vitro.

### LPS/IFN-γ-Induced Intestinal Inflammation Model Recapitulates IBD Pathology

To establish an IBD model, we treated intestinal organoids with lipopolysaccharide (LPS) and interferon-gamma (IFN-γ) to simulate the inflammatory milieu characteristic of IBD. In the inflammation model group, the epithelial architecture was markedly disrupted, with dead cells shed into the organoid lumens — a phenotype consistent with the epithelial denudation observed in active IBD lesions (Figure 5A).

**Figure 5.**
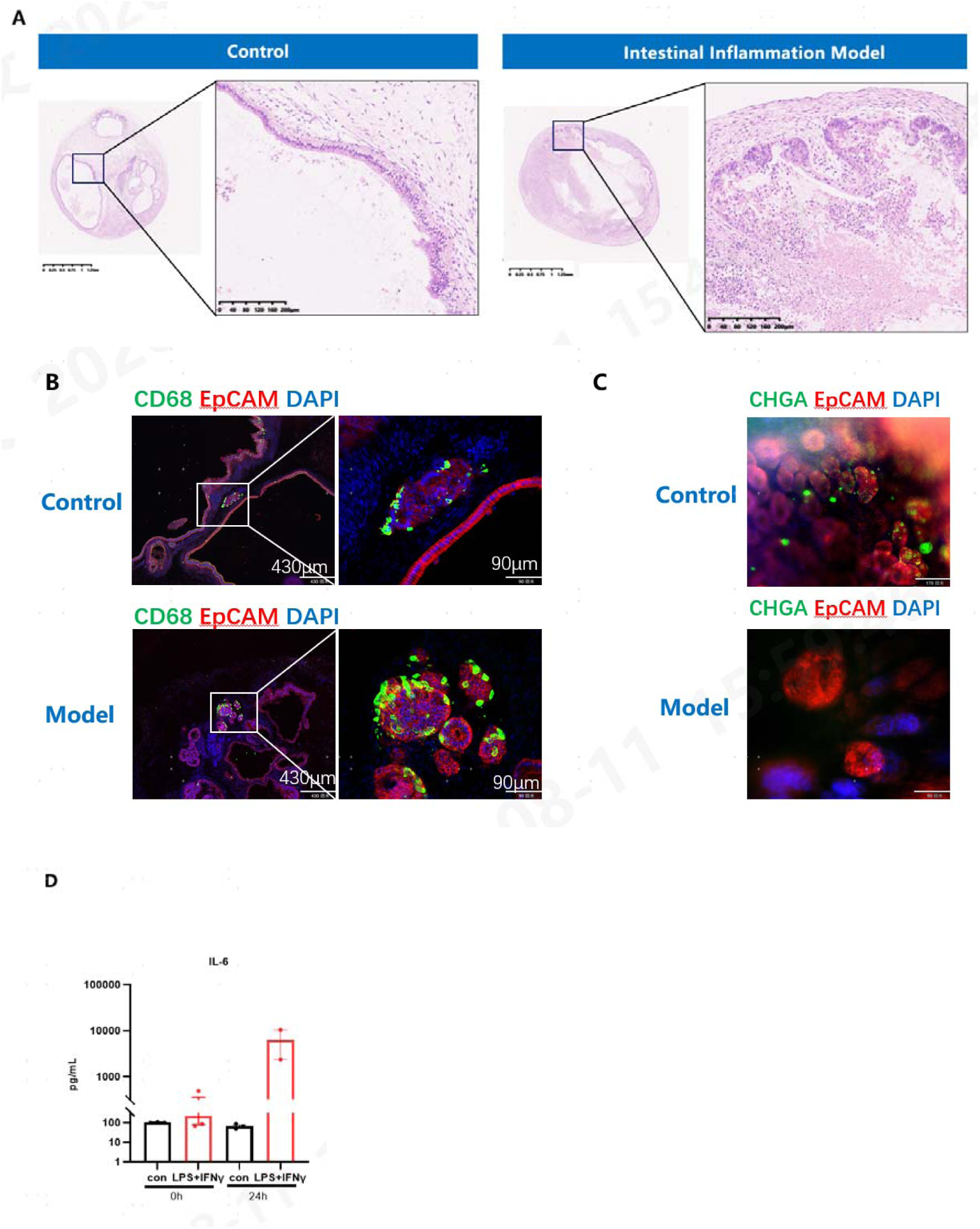
LPS/IFN-γ-induced intestinal inflammation model. (A) H&E images of control and inflammation model organoids, showing epithelial disruption and dead cell shedding into the lumen in the model group. Scale bars: 1.25 mm (left), 200 μm (right). (B) Immunofluorescence for CD68 (immune cells) and EPCAM (epithelial cells) in control vs. model organoids at 0 h and 24 h. Scale bars: 430 μm (left), 90 μm (right). (C) Immunofluorescence for CHGA (neuroendocrine cells) and EPCAM in control vs. model organoids, showing reduced epithelial and neuroendocrine cells in the model group. Scale bar: 90 μm. (D) IL-6 levels in culture supernatants at 0 h and 24 h post-stimulation (control vs. LPS+IFN-γ). Data are mean ± SD; **p < 0.01, ***p < 0.001.

Immunofluorescence analysis revealed significant changes in cellular composition following inflammatory stimulation. CD68+ immune cells showed increased abundance and activation in the model group compared to controls, indicating immune cell infiltration and macrophage activation (Figure 5B). Concurrently, EPCAM+ epithelial cells were markedly reduced in the inflammation model, reflecting epithelial damage and loss (Figure 5C). CHGA+ neuroendocrine cells were also diminished in the model group, suggesting that inflammation affects specialized epithelial lineages beyond absorptive cells (Figure 5C).

Cytokine analysis demonstrated that IL-6 levels were significantly elevated at 24 hours post-stimulation in the LPS + IFN-γ group compared to unstimulated controls (Figure 5D), confirming the induction of a pro-inflammatory cytokine response. The time-dependent increase in IL-6 is consistent with the acute inflammatory phase of IBD.

### Transcriptomic Analysis Confirms IBD-Related Pathway Activation

To gain mechanistic insight into the inflammatory response, we performed bulk RNA sequencing (RNA-seq) on control and LPS/IFN-γ-stimulated intestinal organoids. Differential expression analysis identified numerous differentially expressed genes (DEGs), with Gene Ontology (GO) enrichment analysis revealing the most significantly altered biological processes (Figure 6A).

**Figure 6.**
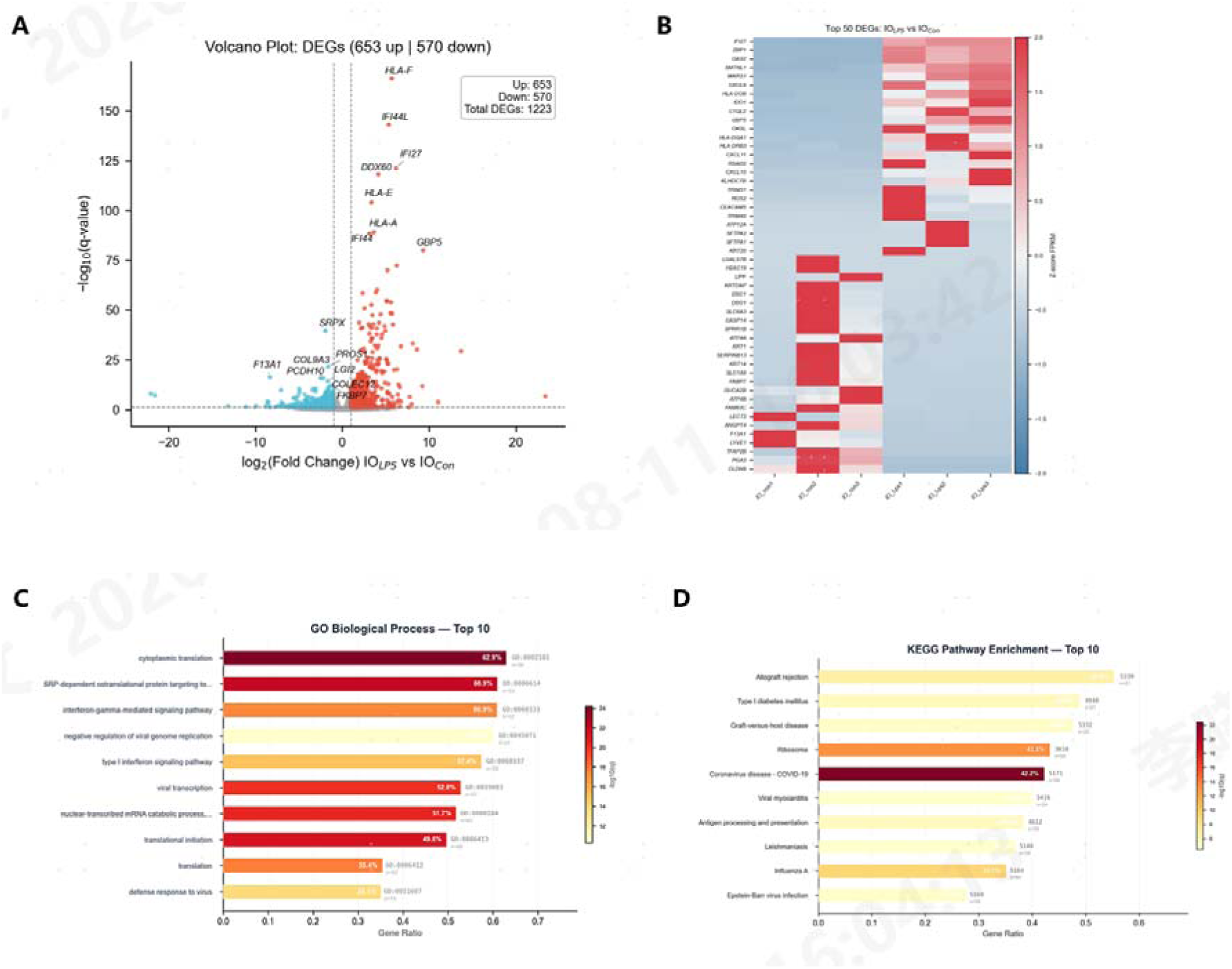
Transcriptomic analysis of the intestinal inflammation model. (A)Volcano plot of differentially expressed genes (DEGs) between control and LPS/IFN-γ-treated organoids (B) Heatmap of TOP50 differentially expressed genes (C) Gene Ontology (GO) enrichment analysis of differentially expressed genes (DEGs) between control and LPS/IFN-γ-treated organoids. Top enriched biological processes include cytoplasmic translation, SRP-dependent co-translational protein targeting, translational initiation, antigen processing, and IFN-gamma response. (D) KEGG enrichment analysis of differentially expressed genes (DEGs) between control and LPS/IFN-γ-treated organoids.

The top enriched terms included cytoplasmic translation, SRP-dependent co-translational protein targeting, and translational initiation, reflecting a massive reprogramming of protein synthesis machinery during inflammation. Immune-related terms, including antigen processing and presentation and IFN-gamma response, were also highly enriched, consistent with the pathological mechanisms of IBD.

IFN-γ Response Pathway Confirmation: The model group showed significant upregulation of a comprehensive set of interferon-stimulated genes (ISGs), confirming the activation of the IFN-γ signaling axis (Figure 6B).

NF-κB Pathway Activation: We observed significant upregulation of RELA (p65), a central transcription factor of the NF-κB pathway. Downstream effectors, including IL6, NOS2 (iNOS), and CCL5, were significantly elevated, and IL33 — an alarmin released upon epithelial damage — was induced. These findings are consistent with the canonical NF-κB-driven inflammatory cascade observed in IBD patients (Figure 6B).

JAK2-STAT3 Axis Activation: We confirmed the IL6 → JAK2 → STAT3 positive feedback loop at the transcriptional level. This pathway is of particular translational significance, as it represents the primary target of Tofacitinib, a JAK inhibitor approved for the treatment of ulcerative colitis. The activation of this axis in our model validates its utility for evaluating JAK inhibitor efficacy (Figure 6B).

Multi-Modal Cell Death Activation: A striking finding was the simultaneous activation of multiple programmed cell death pathways. We observed significant upregulation of CASP1 (the executor of pyroptosis), MLKL and ZBP1 (key mediators of necroptosis), and FAS (a death receptor triggering apoptosis). The concurrent activation of pyroptosis, necroptosis, and apoptosis represents the core mechanism of IBD epithelial damage and may explain the severe epithelial denudation observed in the model (Figure 6C).

Immune Recruitment Machinery: We observed significant upregulation of antigen presentation molecules (MHC-II, including HLA-DQA1) and leukocyte adhesion molecules (ICAM1, VCAM1). This indicates active antigen presentation and leukocyte adhesion within the organoids, mimicking the immune cell recruitment cascade that drives chronic intestinal inflammation (Figure 6C).

### Establishment of Intestinal Cancer Models

To evaluate the applicability of the organoid platform for cancer modeling, we established intestinal cancer models at two time points. In the 7-day cancer model, H&E staining revealed elongated, hyperchromatic nuclei that retained polarity and exhibited pseudostratification — features indicative of early neoplastic transformation (Figure 7A). Immunofluorescence staining for Ki67 and CK20 demonstrated abnormal cellular hyperplasia in the cancer model group compared to controls, with increased Ki67+ proliferating cells and disorganized CK20+ epithelial structures (Figure 7C).

**Figure 7.**
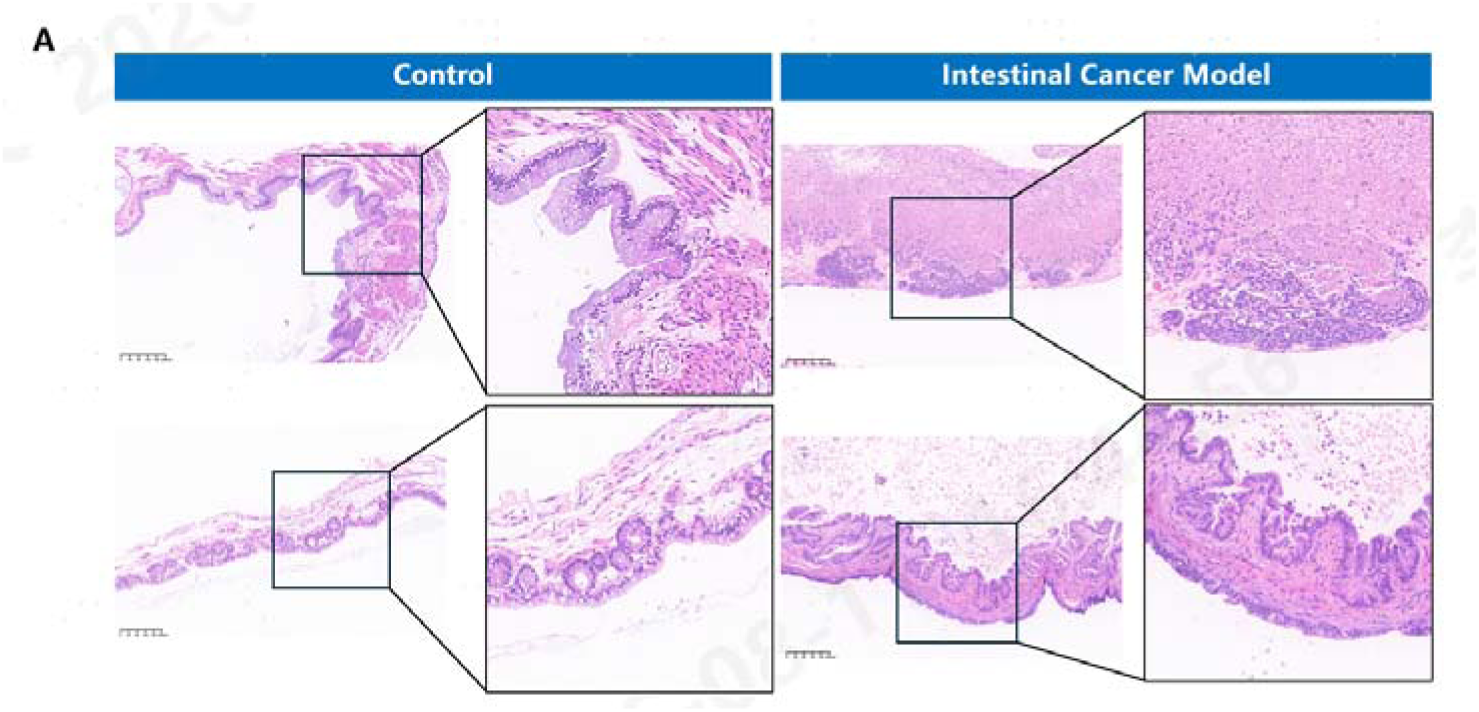
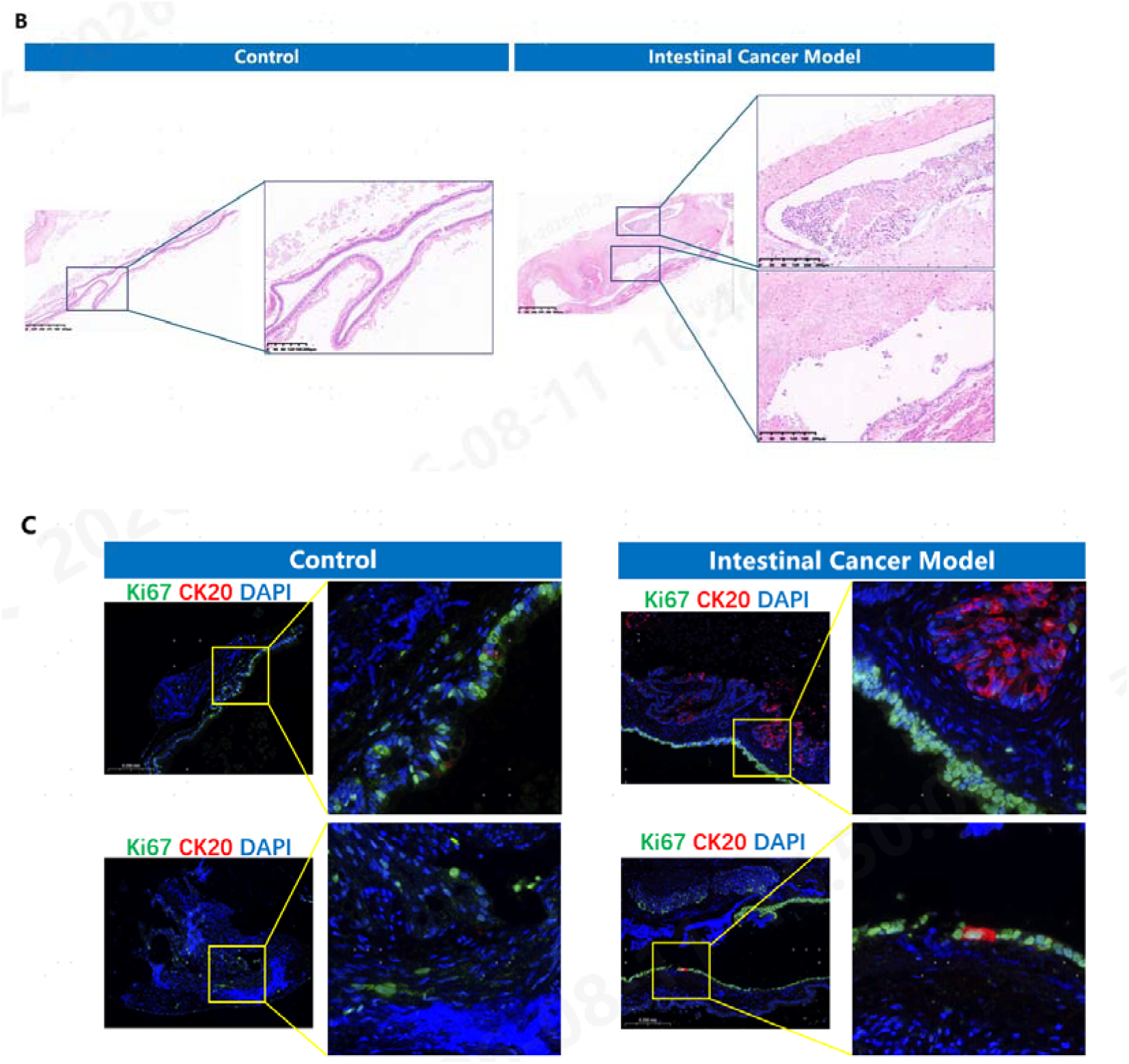
Intestinal cancer models. (A) H&E staining of 7-day cancer model organoids compared to controls, showing elongated, hyperchromatic nuclei with pseudostratification. Scale bars: 200 μm. (B) H&E staining of 21-day cancer model organoids compared to controls, showing advanced neoplastic features. Scale bars: 200 μm. (C) Immunofluorescence for Ki67 (proliferation) and CK20 (epithelial differentiation) in 7-day control vs. cancer model organoids, showing abnormal hyperplasia. Scale bars: 90 μm.

In the 21-day cancer model, we observed more advanced phenotypic changes, including further architectural distortion and increased cellular atypia (Figure 7B). The progression from early pseudostratification at 7 days to more pronounced neoplastic features at 21 days suggests that this model can capture the temporal dynamics of intestinal tumorigenesis.

### Probiotic–Intestinal Organoid Co-Culture Model

To demonstrate the utility of the platform for evaluating microbe-host interactions, we developed a probiotic–intestinal organoid co-culture system. We microinjected fluorescently labeled probiotics into the lumens of intestinal organoids, allowing direct visualization of bacterial localization within the organoid lumen (Figure 8A).

**Figure 8.**
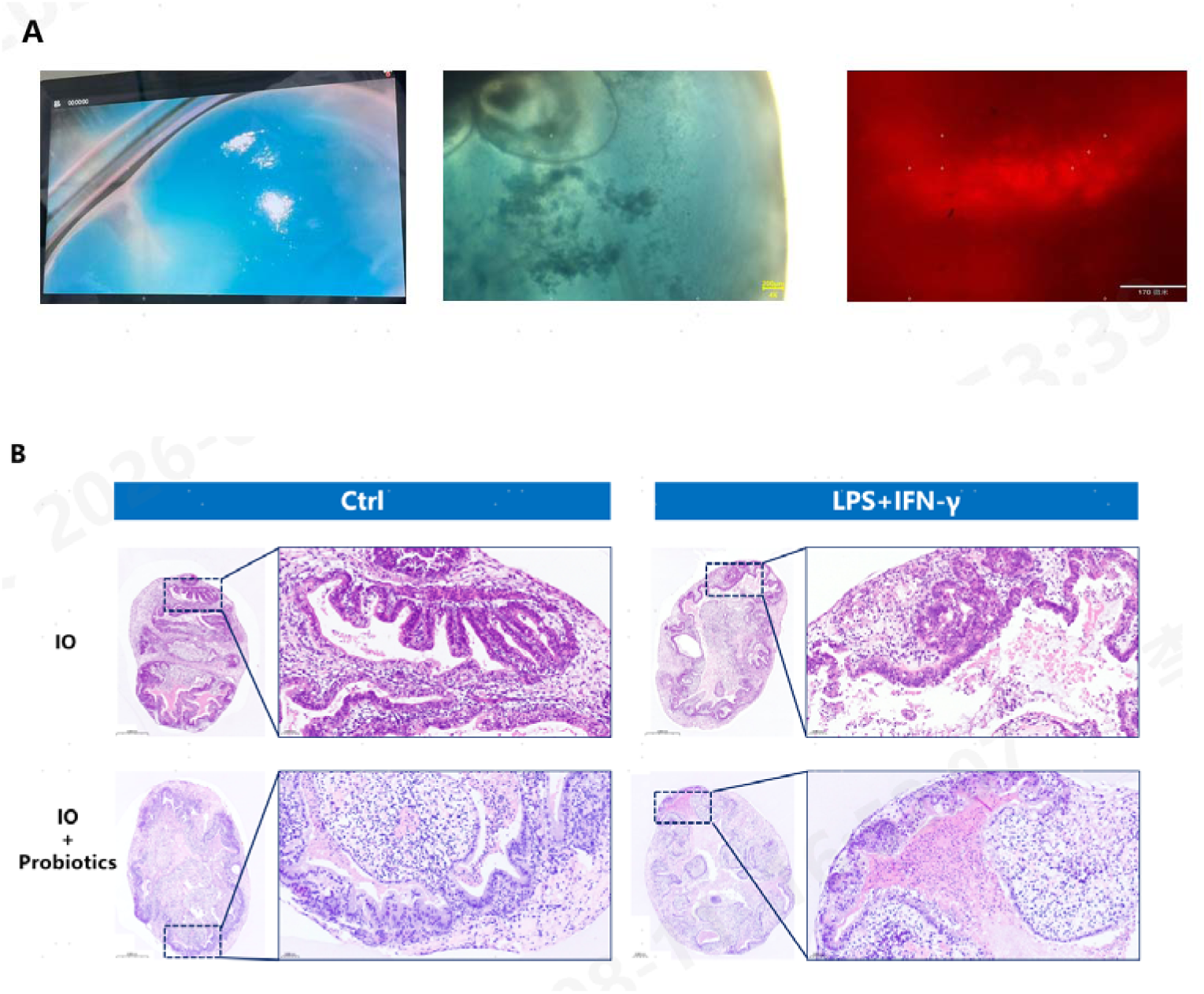
Probiotic–intestinal organoid co-culture model. (A) Fluorescence microscopy images showing green fluorescent dye-labeled probiotics microinjected into the lumen of an intestinal organoid (IO). Scale bar: 500 μm. (B) Comparison of inflammatory markers across four conditions: control, LPS+IFN-γ, IO-52D + probiotics, and LPS+IFN-γ + probiotics. Probiotic co-culture reduces inflammatory markers in the LPS/IFN-γ-treated group.

Using the IO-52D organoid line, we induced inflammation with LPS + IFN-γ and evaluated the anti-inflammatory effects of probiotic co-culture. The probiotic-treated group demonstrated reduced inflammatory markers compared to the LPS + IFN-γ-only group, indicating that the co-culture system can detect and quantify the immunomodulatory effects of probiotics (Figure 8B). This establishes the platform as a functional readout system for evaluating beneficial microorganisms and their impact on intestinal inflammation.

## Discussion

In this study, we describe the development and characterization of Centimeter-Scale, purely 3D self-organized human intestinal organoids derived from iPSCs, and demonstrate their application as a multicellular platform for modeling IBD, colorectal cancer, and host-microbe interactions. Our findings highlight several key advances over conventional intestinal organoid models.

### Technical advantages of Centimeter-Scale 3D self-organization

Traditional intestinal organoids, whether derived from adult stem cells or iPSCs, are typically cultured as small spheroids in Matrigel domes and are predominantly epithelial in composition. While these models have been invaluable for studying epithelial biology, they are intrinsically limited in their ability to model diseases that involve multi-tissue interactions. Our purely 3D self-organization approach yields organoids that reach centimeter-scale dimensions and spontaneously develop multiple tissue lineages without exogenous stromal co-culture. Relative to conventional systems, our platform offers four distinct advantages. First, the neuro-muscle lineages are generated autonomously in vitro, mature to form functional neuromuscular junctions, and produce macroscopically visible peristaltic-like contractions that mimic intestinal motility without xenotransplantation. Second, the differentiation is performed entirely in 3D, with straightforward handling and high success rates. Third, the protocol is completely free of Matrigel, allowing culture in bioreactors for high-throughput production under fully defined and stable conditions. Fourth, the organoids reach the largest reported size for intestinal organoids and remain robust, with survival and peristalsis maintained for over 120 days in culture. The emergence of rhythmic peristaltic-like contractions at day 100+ represents a functional milestone rarely achieved in conventional organoid systems. This motility reflects the maturation of functional smooth muscle-neuron units, which are essential for modeling motility disorders and drug effects on intestinal contractility.

### Multicellular complexity enables physiologically relevant disease modeling

The single-cell transcriptomic analysis at day 115 revealed 12 cell subtypes across four major lineages, including multiple fibroblast subtypes with distinct signaling roles. The presence of crypt fibroblasts expressing RSPO3 and WNT2 — key niche factors supporting epithelial stemness — indicates the establishment of authentic epithelial-mesenchymal crosstalk. The presence of ENS-associated fibroblasts (SOX11+) and neuronal cells further demonstrates the formation of enteric nervous system components, increasingly recognized as critical players in IBD pathogenesis through neuro-immune interactions (Margolis et al., 2016). Furthermore, the endogenous presence of CD68+ immune cells eliminates the need for exogenous immune cell addition, a significant advantage over co-culture systems that require careful optimization of immune cell ratios and activation states.

### IBD model recapitulates clinical pathology at multiple levels

The LPS/IFN-γ-induced inflammation model reproduced key features of IBD at morphological, cellular, and molecular levels. The epithelial disruption and dead cell shedding into lumens mirror the epithelial denudation seen in active IBD. The transcriptomic analysis revealed activation of canonical IBD pathways — NF-κB, JAK2-STAT3, and IFN-γ response — that are directly targeted by currently approved therapies (e.g., anti-TNF biologics, Tofacitinib). The discovery of concurrent pyroptosis, necroptosis, and apoptosis activation is particularly noteworthy, as this multi-modal cell death phenotype represents a core mechanism of IBD epithelial damage that has been difficult to model in epithelial-only systems. The upregulation of antigen presentation machinery (MHC-II) and leukocyte adhesion molecules (ICAM1, VCAM1) further demonstrates that the model captures the immune recruitment cascade central to chronic intestinal inflammation.

### Cancer modeling and the inflammation-cancer axis

The intestinal cancer models at 7 and 21 days demonstrated progressive neoplastic features, from early pseudostratification to advanced architectural distortion. The ability to model both inflammation and cancer within the same organoid platform opens opportunities for studying the "inflammation-to-cancer" transition — a clinically important sequence in IBD-associated colorectal cancer (IBD-CRC) that is difficult to recapitulate in conventional models. Future studies could combine the IBD and cancer models to investigate how chronic inflammation promotes malignant transformation in a multicellular context.

### Probiotic co-culture system

The microinjection-based probiotic delivery system enables direct assessment of microbial effects on the intestinal epithelium and immune compartment within a controlled 3D environment. The observed anti-inflammatory effects of probiotics in the LPS/IFN-γ model validate this system as a functional assay for evaluating beneficial microorganisms, with potential applications in probiotic screening and microbiome-targeted therapy development.

### Limitations and future directions

This study has several limitations that should be addressed in future work. First, the current data are primarily derived from a single iPSC line; validation across multiple donor lines and disease-specific iPSCs (e.g., from IBD or CRC patients) would strengthen the generalizability of the findings. Second, although we observed CD31+ vascular-like structures, the organoids lack perfusable vasculature, which limits long-term culture and modeling of vascular-related pathology. Integration with vascularization strategies, such as co-culture with endothelial cells or organ-on-chip platforms, could address this limitation. Third, the probiotic co-culture system currently involves single-strain delivery; expanding to complex microbial communities would better recapitulate the native microbiome-host interaction. Finally, the inflammatory and cancer models would benefit from centimeter-scale validation with quantitative histomorphometric analysis and correlation with patient-derived data.

In conclusion, Centimeter-Scale, multicellular intestinal organoids derived from human iPSCs provide a physiologically relevant platform that overcomes key limitations of conventional epithelial-only organoid models. The combination of multilineage cellular diversity, functional maturation, and demonstrated applicability to IBD, CRC, and host-microbe interaction modeling positions this platform as a valuable tool for intestinal disease research, drug discovery, and personalized medicine.

## Methods

### iPSC Culture and Intestinal Organoid Differentiation

Human iPSCs were maintained in mTeSR Plus medium (STEMCELL Technologies) on Matrigel (Corning)-coated plates at 37°C, 5% COO. Differentiation was initiated at ∼80% confluence. The differentiation protocol followed a modified directed differentiation approach: definitive endoderm induction (days 0–3), hindgut specification (days 3–7), and 3D intestinal organoid formation (days 7–9). From day 9 onward, organoids were maintained in proprietary Intestinal Organoid Maintenance Medium (MIDIO-001, Neocellmed) with medium changes every 3–4 days. Human iPSCs are seeded at 9,000 cells per well in 96-well U-bottom ultra-low attachment plates with 100 µL of KIDIO-001-A medium (Day 0) and cultured for 4 days with half-medium changes every 48 hours to form compact, round embryoid bodies (EBs) of 200–300 µm diameter. On Day 4, the medium is switched to KIDIO-001-B and cultured for another 5 days (until Day 9) with medium changes every 48 hours, during which EBs enlarge and show polarization. On Day 9, EBs are carefully transferred to Erlenmeyer flasks containing KIDIO-001-C medium and placed on an orbital shaker at 70 rpm for 11 days with complete medium changes every 48 hours, resulting in cystic structures up to 3–5 mm. From Day 20, the medium is changed to KIDIO-001-D maintenance medium and culture continues under the same shaking conditions with complete medium changes every 48 hours until Day 42 or beyond, yielding organoids of 1–3 cm diameter with crypt-like domains, goblet cells, and smooth muscle-like layers.

For long-term maintenance or disease modeling (e.g., IBD, intestinal cancer), the medium is switched to KIDIO-001-M from Day 42 onwards, with changes every 3–4 days, allowing organoids to be maintained for over 120 days with functional activity; after Day 100, rhythmic peristaltic-like contractions may be observed

### Histology and Immunofluorescence

For H&E staining, organoids were fixed in 4% paraformaldehyde (PFA) for 2 hours at 4°C, embedded in paraffin, sectioned at 5 μm, and stained with hematoxylin and eosin following standard protocols. For immunofluorescence on sections, paraffin sections were deparaffinized, subjected to antigen retrieval in citrate buffer (pH 6.0), blocked with 5% normal donkey serum, and incubated with primary antibodies overnight at 4°C. The following primary antibodies were used: anti-EPCAM (1:200, Abcam), anti-CD68 (1:100, Cell Signaling Technology), anti-CHGA (1:200, Abcam), anti-α-SMA (1:400, Sigma), anti-TUJ1 (1:500, BioLegend), anti-CD31 (1:100, Dako), anti-Ki67 (1:200, Abcam), and anti-CK20 (1:200, Abcam). Secondary antibodies were Alexa Fluor-conjugated (Invitrogen, 1:500). Nuclei were counterstained with DAPI. For whole-mount immunofluorescence, intact organoids were fixed in 4% PFA, permeabilized with 0.5% Triton X-100, blocked, and incubated with primary and secondary antibodies sequentially. Imaging was performed on a confocal laser scanning microscope (Leica SP8 or Zeiss LSM 880).

### Single-Cell RNA Sequencing

Day 115 intestinal organoids were dissociated into single cells using Accutase (STEMCELL Technologies) at 37°C for 20 minutes with intermittent pipetting. Cell viability was assessed by trypan blue exclusion (>85% viability required). Single-cell libraries were prepared using the 10x Genomics Chromium platform according to the manufacturer’s protocol. Sequencing was performed on an Illumina NovaSeq 6000. Raw sequencing data were processed using Cell Ranger (10x Genomics) for alignment and UMI counting against the human reference genome (GRCh38). Downstream analysis was performed using Seurat (v4). Cells with <200 genes, >20% mitochondrial reads, or doublet scores above threshold were removed. Data were normalized using SCTransform, and dimensionality reduction was performed using PCA followed by UMAP. Clustering was performed using the Louvain algorithm. Cell types were annotated based on canonical marker gene expression and cross-referenced with published intestinal cell atlases.

### Intestinal Inflammation Model

Intestinal organoids (day 52+) were treated with LPS (100 ng/mL, E. coli O111:B4, Sigma) and recombinant human IFN-γ (50 ng/mL, PeproTech) for 24 hours. Control organoids received vehicle only. Morphological changes were monitored by brightfield microscopy. After treatment, organoids were harvested for histology, immunofluorescence, RNA-seq, and cytokine analysis.

### Cytokine Measurement

IL-6 levels in organoid culture supernatants were measured at 0 and 24 hours post-stimulation using a human IL-6 ELISA kit (R&D Systems) according to the manufacturer’s instructions. Absorbance was read at 450 nm on a microplate reader.

### Bulk RNA Sequencing and Analysis

Total RNA was extracted from control and LPS/IFN-γ-treated organoids using TRIzol (Invitrogen). RNA quality was assessed using an Agilent Bioanalyzer (RIN > 7.0). Libraries were prepared using the NEBNext Ultra II RNA Library Prep Kit and sequenced on an Illumina NovaSeq 6000 (paired-end 150 bp). Reads were aligned to the human reference genome (GRCh38) using STAR. Differential expression analysis was performed using DESeq2 (|logOFC| > 1, adjusted p < 0.05). Gene Ontology enrichment analysis was performed using clusterProfiler (R package). Heatmaps were generated using pheatmap.

### Intestinal Cancer Model

Intestinal organoids were treated with a chemical carcinogen cocktail [specific agents and concentrations to be detailed] for 7 days (7D model) or 21 days (21D model). Control organoids received vehicle. Tissues were processed for H&E staining and immunofluorescence as described above.

### Probiotic–Organoid Co-Culture

Probiotic strains [strain identity to be detailed] were labeled with a green fluorescent dye (CFSE or FITC) according to the manufacturer’s protocol. Labeled probiotics were microinjected into the lumens of intestinal organoids (IO-52D) using a microinjector (Eppendorf FemtoJet). For the anti-inflammatory evaluation, probiotic-injected organoids were simultaneously treated with LPS (100 ng/mL) and IFN-γ (50 ng/mL). Inflammatory markers were assessed at 24 hours and compared to LPS/IFN-γ-only and untreated control groups.

### Statistical Analysis

All quantitative data are presented as mean ± standard deviation (SD) from at least three biological replicates unless otherwise stated. Statistical comparisons between two groups were performed using Student’s two-tailed t-test. Multiple group comparisons were performed using one-way ANOVA with Tukey’s post hoc test. p < 0.05 was considered statistically significant. All analyses were performed using GraphPad Prism (v9.0) or R (v4.2.0).

### Data Availability

Single-cell RNA-seq and bulk RNA-seq data have been deposited in the Gene Expression Omnibus (GEO) under accession number [to be submitted upon acceptance]. All other data supporting the findings of this study are available from the corresponding author upon reasonable request.

## Supporting information

Video of the peristalsis of intestinal organoids

## Acknowledgments

We thank all members of the research and development team at Shanghai Neocellmed Co., Ltd. for their technical assistance and discussions. We acknowledge the use of core facilities for sequencing and microscopy.

## Author Contributions

[To be filled: Conceptualization, Methodology, Investigation, Data Analysis, Writing – Original Draft, Writing – Review & Editing, Supervision, Funding Acquisition]

## Supplementary Information

**Figure S1. Supplementary Figure S1**

Figure S1. The bright-field video shows the folded epithelial structure and peristalsis of the intestinal organoid after about 100 days.

**Figure S2. Supplementary Figure S2**

Figure S2. Video of the peristalsis of a 10 cm intestinal organoid

**Figure S3. Supplementary Figure S3**

Figure S3. Video of the peristalsis of 3-5 cm intestinal organoids

